# Animal complexity defines the functionality of synonymous mutations through mediating microRNA regulations

**DOI:** 10.1101/2022.08.30.505799

**Authors:** Chunmei Cui, Xiao Lin, Qinghua Cui

**Affiliations:** Department of Biomedical Informatics, MOE Key Lab of Cardiovascular Sciences, School of Basic Medical Sciences, Peking University, 38 Xueyuan Rd, Beijing, 100191, China

## Abstract

Most recently, Shen et al. revealed that most synonymous mutations (SMs) in yeast genes are strongly non-neutral, however, which was soon challenged by Kruglyak et al. who find no evidence for deleterious effect on fitness of SMs in yeast genes. We previously revealed that SMs in numerous human genes show functionality through changing miRNA regulations. Given that higher animals show higher complexity of miRNA regulations, here we present a hypothesis that SMs in lower animal genes show less functionality and vice versa. To confirm this, we comprehensively investigated the SMs changing miRNA regulations in yeast, *C.elegans, Drosophila*, mouse, and human. As expected, the result showed that the proportion of SMs affecting gene functionality through altering miRNA regulations is about 0%, 14%, 16%, 51%, and 60% in yeast, *C. elegans, Drosophila*, mouse, and human, respectively, suggesting that the complexity of animals indeed determines the functionality of SMs through mediating miRNA regulations. Finally, this study supports no evidence that most SMs in yeast genes are deleterious.

## Introduction

Synonymous mutations (SMs) represent one of the most common genetic variations and are traditionally considered to be neutral in evolution and thus not functional as they do not change protein sequences and structures. However, growing evidences have revealed that SMs could affect the functionality of gene products by diverse mechanisms, for instance, regulatory binding-sites for transcription factors and microRNAs (miRNAs), precursor mRNA splicing, and mRNA stability and translation etc. ^1–3^. Most recently, Shen et al presented that most SMs in 21 yeast genes are strongly non-neutral and deleterious ^4^, however, Kruglyak et al argued that nearly all SMs have no detectable influence on the fitness ^5^, which has sparked controversy on whether SMs are deleterious/functional or not again. We previously revealed that nearly half of human SMs could affect gene functionality through disrupting miRNA regulation ^6^. Moreover, it is well known that miRNA regulation shows significantly greater complexity in higher animals ^7^. For example, no miRNAs are found in yeast but 2656 miRNAs have been identified in human. Given the above observations, here we propose a hypothesis that SMs in lower animal (e.g. yeast) genes show less functionality and vice versa. To confirm this hypothesis, we comprehensively investigated the SMs changing miRNA regulations in five animals, including yeast, *C. elegans*, *Drosophila*, mouse, and human. As a result, the proportions of SMs changing miRNA regulations in yeast, C.elegans, Drosophila, mouse, and human are around 0%, 14%, 16%, 51%, and 60%, respectively, suggesting that SMs show no functionality in yeast genes but show significant functionality in higher animals, such as human and mouse.

## Method

The single nucleotide polymorphism (SNP) data of human, mouse, and *Drosophila*, were collected from dbSNP repository ^8^. Notably, here we only fetched common SNPs in human for further analysis. And, the SNPs occurring in *C. elegans* genome were obtained in WormBase ^9^. Meanwhile, the genome annotation info and gene sequences were downloaded from NCBI database and WormBase, respectively. We searched for the SNPs located in gene CDS regions by mapping SNPs’ position to the genome and retained SMs for further analysis. The miRNA sequences were obtained from miRBase ^10^.

To examine SMs that could alter miRNA-mediated gene regulation, TargetScan^11^ and miRanda^12^, two of the most popular tools for predicting miRNA targets, were employed. To be specific, for each SM, we selected the 15nt flanking sequences around mutation sites and generated two sequences with the reference and alternative allele at the center, then two sequences were separately predicted the miRNA binding sites for detecting the change of regulating miRNA after mutation. Besides, we adopted the intersection result of TargetScan and miRanda for each sequence.

## Results

Firstly, we found 154,714, 301,236, 390,895, and 141,433 SMs in human, mouse, *Drosophila* and *C. elegans*, respectively. Subsequently, we identified the SMs with changing miRNA-mediated gene regulations in the four animals. And then we calculated the proportion of SMs altering miRNA regulations in the four animals (*C. elegens, Drosophila*, mouse, and human). In addition, because no miRNAs are found in yeast, the proportion of SMs altering miRNA regulations in yeast will be 0. As a result, the proportions of SMs altering miRNA regulations are 0%, 14.11%, 15.73%, 50.67%, and 60.06% in yeast, *C. elegans, Drosophila*, mouse, and human, respectively (Figure 1), suggesting SMs in higher animal genes have greater functionality. For example, the proportions of SMs altering miRNA regulation in human (60.06%) and mouse (50.67%) is significantly greater than (P = 0, Chi-Square test) that in *Drosophila* (15.73%) and *C. elegans* (14.11%). The result supports our hypothesis that SMs in lower animal (e.g. yeast) genes show less functionality and vice versa. Specifically, our study supports no evidence that SMs in yeast genes are mostly deleterious.

**Figure 1.**
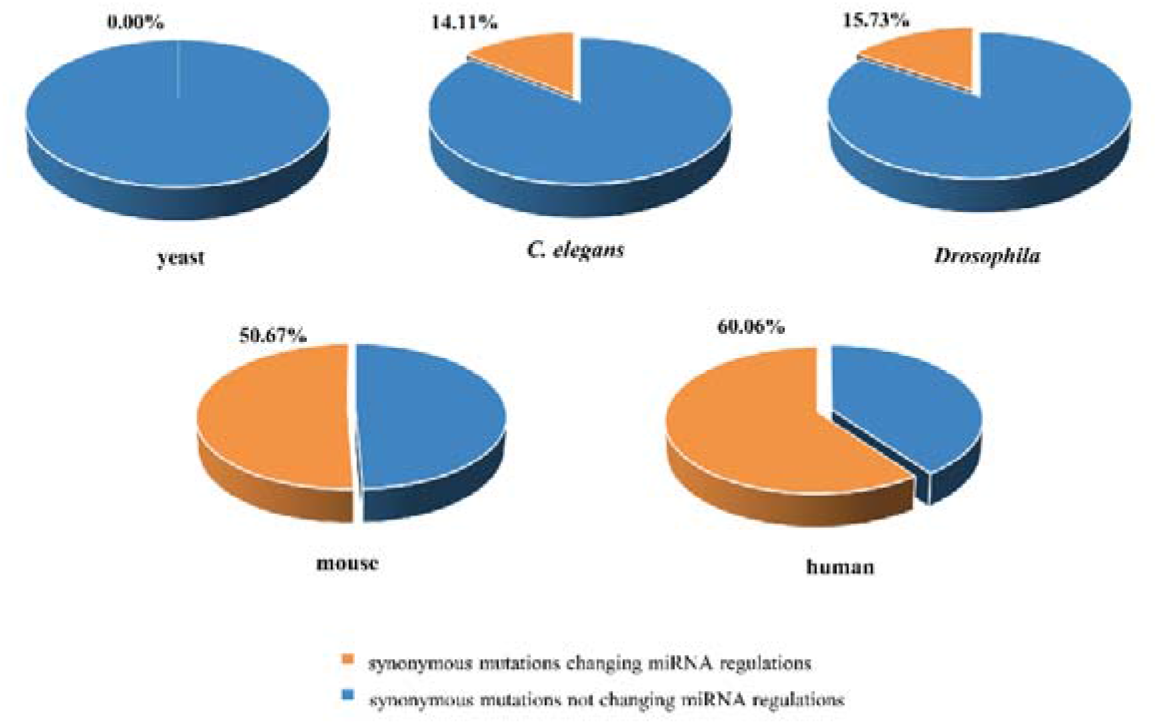
The proportions of synonymous mutations changing miRNA regulation in yeast, *C. elegans, Drosophila*, mouse, and human.

## Discussion

In summary, we found the proportion of SMs with changing miRNA regulation monotonically increases as the species becomes more complex, which supports our hypothesis that SMs in lower animal genes show less functionality and vice versa. Given that there is no miRNA found in yeast, our model suggests no evidence that synonymous mutations in yeast genes are mostly deleterious.

## Acknowledgment

This work has been supported by the grant from the Natural Science Foundation of China (62025102, 81921001, 81970440).

## Conflict of Interest

none declared.

## Notes

### Competing Interest Statement

The authors have declared no competing interest.

## References

1 Sauna, Z. E. & Kimchi-Sarfaty, C. Understanding the contribution of synonymous mutations to human disease. Nature Reviews Genetics 12, 683–691, doi:10.1038/nrg3051 (2011).

2 Plotkin, J. B. & Kudla, G. Synonymous but not the same: the causes and consequences of codon bias. Nature Reviews Genetics 12, 32–42, doi:10.1038/nrg2899 (2011).

3 Hunt, R., Sauna, Z. E., Ambudkar, S. V., Gottesman, M. M. & Kimchi-Sarfaty, C. Single Nucleotide Polymorphisms: Methods and Protocols (ed Anton A. Komar) 23–39 (Humana Press, 2009).

4 Shen, X., Song, S., Li, C. & Zhang, J. Synonymous mutations in representative yeast genes are mostly strongly non-neutral. Nature 606, 725–731, doi:10.1038/s41586-022-04823-w (2022).

5 Kruglyak, L. et al. No evidence that synonymous mutations in yeast genes are mostly deleterious. bioRxiv. Preprint, doi:10.1101/2022.07.14.500130 (2022).

6 Wang, Y., Qiu, C. & Cui, Q. A Large-Scale Analysis of the Relationship of Synonymous SNPs Changing MicroRNA Regulation with Functionality and Disease. International journal of molecular sciences 16, 23545–23555, doi:10.3390/ijms161023545 (2015).

7 Mao, X., Li, L. & Cao, Y. Evolutionary comparisons of miRNA regulation system in six model organisms. Genetica 142, 109–118, doi:10.1007/s10709-014-9758-5 (2014).

8 Sayers, E. W. et al. Database resources of the National Center for Biotechnology Information. Nucleic acids research 49, D10–d17, doi:10.1093/nar/gkaa892 (2021).

9 Harris, T. W. et al. WormBase: a modern Model Organism Information Resource. Nucleic acids research 48, D762–d767, doi:10.1093/nar/gkz920 (2020).

10 Kozomara, A., Birgaoanu, M. & Griffiths-Jones, S. miRBase: from microRNA sequences to function. Nucleic acids research 47, D155–d162, doi:10.1093/nar/gky1141 (2019).

11 Agarwal, V., Bell, G. W., Nam, J. W. & Bartel, D. P. Predicting effective microRNA target sites in mammalian mRNAs. eLife 4, doi:10.7554/eLife.05005 (2015).

12 Betel, D., Koppal, A., Agius, P., Sander, C. & Leslie, C. Comprehensive modeling of microRNA targets predicts functional non-conserved and non-canonical sites. Genome biology 11, R90, doi:10.1186/gb-2010-11-8-r90 (2010).

